# Integration of plant carbohydrate dynamics by Fourier polynomials

**DOI:** 10.1101/2021.03.16.435635

**Authors:** Charlotte Seydel, Julia Biener, Vladimir Brodsky, Svenja Eberlein, Thomas Nägele

## Abstract

Quantification of system dynamics is a central aim of mathematical modelling in biology. Defining experimentally supported functional relationships between molecular entities by mathematical terms enables the application of computational routines to simulate and analyse the underlying molecular system. In many fields of natural sciences and engineering, trigonometric functions are applied to describe oscillatory processes. As biochemical oscillations occur in many aspects of biochemistry and biophysics, Fourier analysis of metabolic functions promises to quantify, describe and analyse metabolism and its reaction towards environmental fluctuations. Here, Fourier polynomials were developed from experimental time-series data and combined with block diagram simulation of plant metabolism to study heat shock response of photosynthetic CO_2_ assimilation and carbohydrate metabolism. Findings suggest that increased capacities of starch biosynthesis stabilize photosynthetic CO_2_ assimilation under transient heat exposure. Among soluble sugars, fructose concentrations were observed to fluctuate least under heat exposure which might be the consequence of high respiration rates under elevated temperature. Finally, Col-0 and two mutants of *Arabidopsis thaliana* with deficiencies in starch and sucrose metabolism were discriminated by fundamental frequencies of Fourier polynomials across different experiments. This suggests balance modelling based on Fourier polynomials as a suitable approach for mathematical analysis of dynamic plant-environment interactions.

**Highlight:** A balance equation model is developed to quantify effects of transient heat exposure on plant carbon assimilation. The model is based on Fourier polynomials for quantitative assessment of system dynamics.

## Introduction

Capturing dynamics in biological systems by mathematical terms is the general aim of biomathematical modelling. Differential equations represent an adequate strategy to describe dynamics over space and time. Ordinary differential equations (ODEs) and partial differential equations (PDEs) have been successfully applied to reveal biological system dynamics and to develop predictive models of growth rates, transcription, translation or metabolic processes (Lopatkin and Collins, 2020). In a metabolic context, ODE models are frequently applied to simulate enzyme kinetic reactions and, by this, to explain dynamics of observed metabolite concentrations. Kinetic models, based on ODEs, have frequently been applied in a broad field of biological research, e.g. in context of metabolic engineering of microbial systems and strain design (Costa *et al.*, 2016), disease research (Ramos *et al.*, 2019), and plant metabolism (Feldman-Salit *et al.*, 2019; Rohwer, 2012; Weiszmann *et al.*, 2018). In contrast to the high diversity of application fields, the principle of ODE kinetic modelling remains conserved: based on genome sequence information or biochemical evidence from literature a biochemical reaction network is established, substrate and product concentrations are quantified, and enzyme kinetics are applied to calculate reaction rates within a metabolic system. Enzyme kinetic parameters, e.g., velocity under substrate saturation (V_max_) or substrate affinity (K_M_) of Michaelis-Menten equations are experimentally determined and used for computationally assisted parameter estimation. A well-defined and experimentally validated kinetic ODE model enables computational simulation and prediction of complex system behaviour. A clear limitation of such an approach, however, is the requirement of kinetic parameters which are (frequently) difficult and/or expensive to quantify. Initiatives like KiMoSys, a public repository of published experimental data, summarise and concentrate data on metabolites, protein abundance and fluxes providing a solid database for model construction and initial development (Mochão *et al.*, 2020). Yet, under highly dynamic conditions, e.g., in a fluctuating environment, it still remains a challenge to resolve system dynamics on an enzyme kinetic level. This is due to the high dynamics of metabolites, transcripts, protein levels and enzyme activities (Espinoza *et al.*, 2010; Nägele and Heyer, 2013).

Although being laborious, development and optimization of ODE kinetic models provides an important and informative mathematical method to study biochemical system behaviour. Simultaneously, however, diverse problems might occur with solving and applying such models due to uncertainties about parameters, model structure, kinetic rate laws or parameter sensitivities (Gutenkunst *et al.*, 2007; Schaber *et al.*, 2009). Depending on the research question focused by a study, an explicit knowledge about enzymatic activities and their dynamics might not be essential to derive a mathematical description of metabolite dynamics. For example, metabolic fluxes might be estimated by tracing labelled atoms or molecules in a metabolic pathway system (Crown *et al.*, 2016). While it is not possible to derive information about single enzyme activities or kinetics from flux estimations, they still provide comprehensive insights into metabolic states and pathway activities, also on a large scale (Basler *et al.*, 2018). Beyond, algorithms and user interfaces have been developed which enable the combination of flux data with relative metabolite levels (Sajitz-Hermstein *et al.*, 2016).

For estimating metabolic functions, i.e., the sum of synthesising and degrading/consuming reactions of metabolite pools, under dynamic environmental conditions we have previously suggested a method for implicit estimation of metabolic functions (Nägele *et al.*, 2016). Similar to flux analysis, dynamics of metabolite concentrations in time-series experiments were used in this approach to derive a time-continuous mathematical function to identify regulatory cascades in metabolic pathways. This approach made use of spline interpolations which were composed by cubic polynomials which were fitted to adjacent pairs of data points in a time-series data set. While such an approach is suitable for accurate data fitting, underlying mathematical functions are mostly not related to biological function and, thus, are less predictive than enzyme kinetic models. In the present study, we applied Fourier analysis to develop a mathematical model based on Fourier polynomials to simulate and analyse dynamics of photosynthesis and carbohydrate metabolism under transient heat exposure. Our approach suggests that diurnal and stress-induced dynamics of plant carbohydrate fixation are reflected by harmonic oscillations which provides a mathematical toolbox to study consequences of a changing environment on plant metabolism.

## Materials and Methods

### Plant cultivation and stress treatment

Plants of *Arabidopsis thaliana*, accession Columbia-0 (Col-0), *spsa1*(AT5G20280, SALK line 148643C) and *pgm1* (AT5G51820; TAIR stock CS3092) were grown on a 1:1 mixture of GS90 soil and vermiculite in a climate chamber under short-day conditions (8 h/16 h light/dark; 100 μmol m−2 s−1; 22 °C/18 °C; 60% relative humidity). The *spsa1* line was confirmed via PCR to be homozygous (data not shown) and activity was found to be decreased to 30-50% of the wildtype Col-0 (Fig. S1). The *pgm1* mutant had a dwarf phenotype and starch content was below the detection limit. After 4 weeks, plants were transferred to a growth cabinet (Conviron®, www.conviron.com) and grown for two further weeks under short day conditions with same settings as in the climate chamber. After 6 weeks, at the day of sampling, temperature in the growth cabinet was kept at 22°C during the first two hours in the light (0h→2h, 22°C). Then, in three independent experiments, temperature was increased to (i) 32°C, (ii) 36°C or (iii) 40°C for a total of 4 hours (2h → 6h, temperature increase). In the control experiment, temperature was set constantly to 22°C. Between 6h and 8h, i.e., until the end of the light period, temperature was set to 22°C in all experiments. A graphical representation of the experimental setup is provided in Fig. S2. Plants were sampled at each timepoint (0h, 2h, 6h, 8h) by cutting the full leaf rosette at the hypocotyl. Samples were immediately frozen in liquid nitrogen and stored at −80°C until further use.

### Quantification of net CO_2_ uptake

Rates of net photosynthesis were recorded within the Conviron® growth cabinet using a WALZ® GFS-3000FL system equipped with measurement head 3010-S (Heinz Walz GmbH, Effeltrich, Germany, https://www.walz.com/). Temperature, light and humidity control of the measurement head was set to follow ambient conditions, i.e., to follow surrounding growth cabinet conditions. A summary of recorded temperature, light and humidity curves is provided in the supplement (Fig. S3). For each genotype and growth condition, i.e., temperature setup, three independent samples were measured.

### Extraction and quantification of carbohydrates

Plant material was ground to a fine powder under constant freezing with liquid nitrogen. The powder was lyophilized for three days and subsequently used for carbohydrate analytics. Starch and soluble carbohydrates were extracted and photometrically determined as described before (Kitashova *et al.*, 2021). Plant powder was incubated with 80% ethanol at 80°C for 30min. After centrifugation, the supernatant was transferred to a new tube and extraction was repeated with the pellet. Supernatants were unified and dried in a desiccator. The starch-containing pellet was hydrolyzed with 0.5 M NaOH for 45min at 95°C. After acclimation to room temperature, 1 M CH_3_COOH was added and the suspension was digested with amyloglucosidase solution, finally releasing glucose moieties from starch granules. Glucose was photometrically determined applying a coupled glucose oxidase/peroxidase/o-dianisidine assay.

Soluble sugars sucrose, glucose and fructose were determined from dried ethanol extracts after dissolving in water. After incubation with 30% KOH at 95 °C, sucrose was quantified using an anthrone assay. Anthrone was dissolved in 14.6 M H_2_SO_4_ (0.14% w/v), incubated with the prepared sample for 30 min at 40 °C and absorbance was determined photometrically at 620 nm. Glucose amount was determined photometrically by a coupled hexokinase/glucose 6-phosphate dehydrogenase assay resulting in NADPH + H^+^ at 340 nm. For fructose quantification, phosphoglucoisomerase was added to the reaction mixture after glucose determination.

### Quantification of SPS activity

Activity of sucrose phosphate synthase (SPS) was determined using the anthrone assay as described previously (Kitashova *et al.*, 2021). In brief, freeze-dried leaf tissue was suspended in extraction buffer containing 50 mM HEPES–KOH (pH 7.5), 10 mM MgCl2, 1 mM EDTA, 2.5 mM DTT, 10% (v/v) glycerol and 0.1% (v/v) Triton-X-100. Following incubation on ice, extracts were incubated for 30 min at 25 °C with a reaction buffer containing 50 mM HEPES–KOH (pH 7.5), 15 mM MgCl2, 2.5 mM DTT, 35 mM UDP-glucose, 35 mM F6P and 140 mM G6P. Reactions were stopped by adding 30%KOH and heating to 95 °C. Sucrose was determined photometrically after incubation with anthrone in H_2_SO_4_.

### Statistics, data fitting and modelling

Statistical analysis was performed in R and R Studio (www.r-project.org) (R Core Team, 2019). Fourier series fitting was done within MATLAB® (www.themathworks.com), and block diagram models were created in Simulink® (www.themathworks.com).

## Results

### A block diagram model based on Fourier polynomials for carbon balancing of plant metabolism

A block diagram model of the central carbohydrate metabolism of plants was developed to integrate experimental data on net photosynthesis (NPS), starch and sugar metabolism (Fig. 1). The net carbon input block, i.e., NPS block, represented a Fourier polynomial describing NPS dynamics depending on genotypes and environments (see Fig. 2, solid lines). This input flux was multiplied by the stoichiometric factor 1/6 to enable quantitative summation with starch and sugar fluxes (unit: μmol C6 h^−1^ gDW^−1^). Carbon balance equation 1 (BE_1_) comprised the summation of NPS rates and negative starch rates, balance equation 2 (BE_2_) additionally comprised summation of negative sugar rates (Fig. 1). As a result, BE_1_ revealed the net carbon flux (in C6 equivalents per hour and gram dry weight) which was left from photosynthetically assimilated CO_2_ after starch synthesis, e.g., for sugar biosynthesis or biomass production. Further, BE_2_ revealed residual net carbon flux after additional sugar biosynthesis.

**Figure 1.**
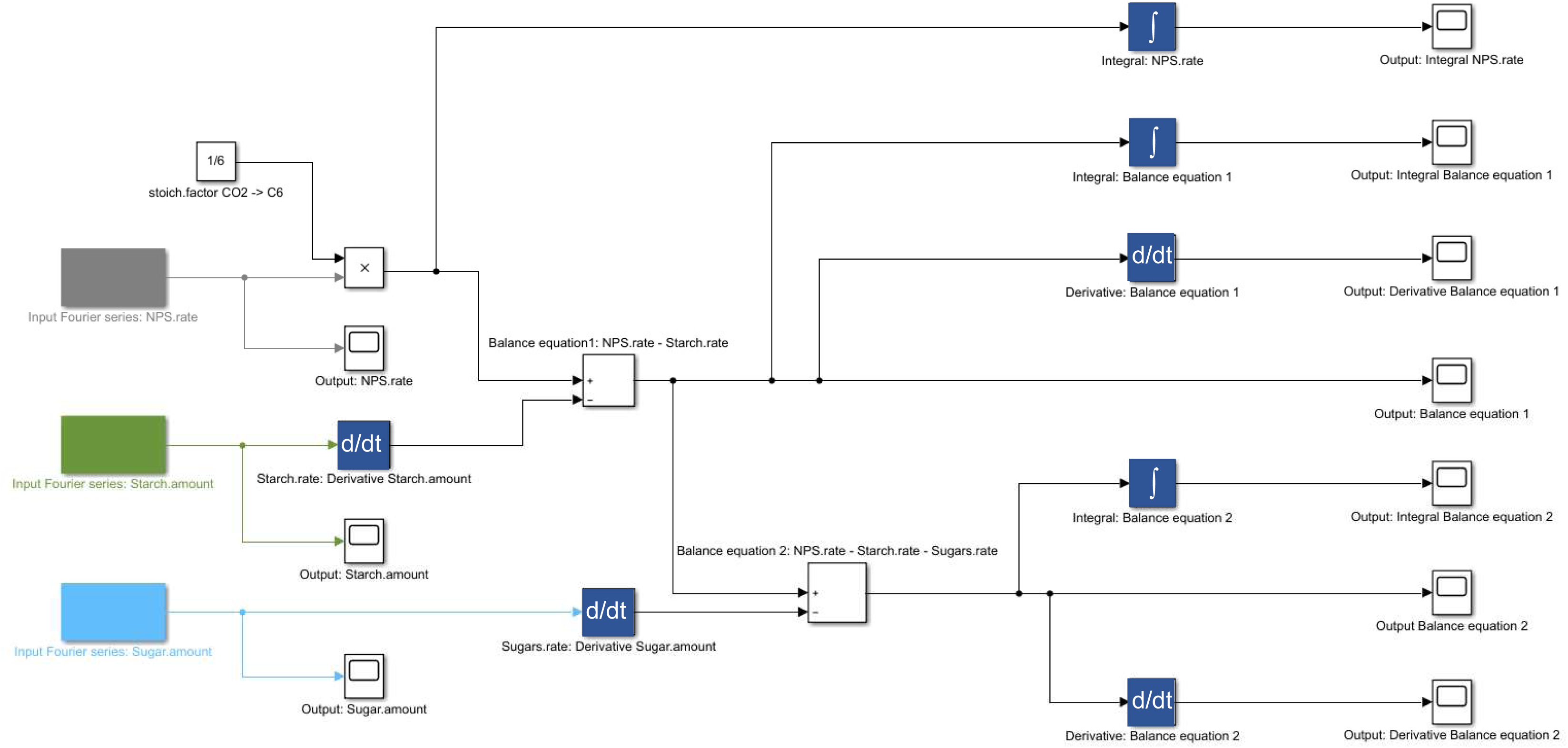
Block diagram applied for Fourier polynomial balance modelling. Input functions are marked in grey (NPS), green (starch amount) and light blue (sugar amount) coloured blocks (left side). Arrows indicate the direction of flux and connect input blocks via multiplication (‘x’) summation (‘+/−‘), differentiation (‘d/dt’) and integration (‘∫’) with output blocks.

**Figure 2.**
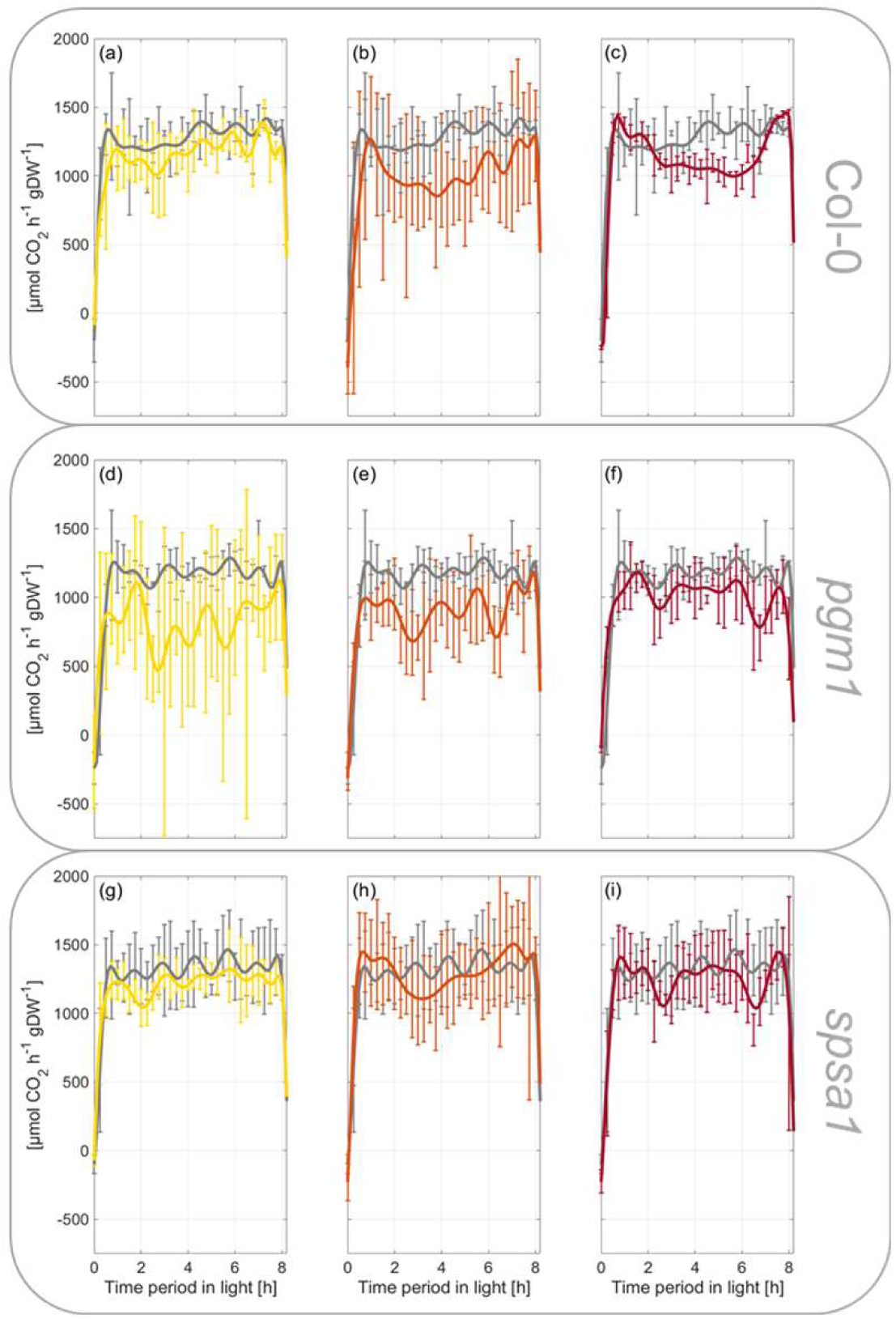
Rates of net CO_2_ uptake during short day transient heat exposure. Error bars represent mean values +/− SD of experimental data (n=3), lines represent Fourier series fits. (a) – (c): Col-0, (d) – (f): *pgm1*, (g) – (i): *spsa1*. Grey lines: 22°C experiment; yellow lines: 32°C experiment; orange lines: 36°C experiment; red lines: 40°C experiment. Temperature was set to 22°C between 0h-2h and 6h-8h. Temperature was transiently increased between 2h and 6h. Temperature curves recorded during the experiments are illustrated in Figure S3a. A summary of Fourier polynomial coefficients is provided in the supplements (Table S1).

Rates of net starch and sugar biosynthesis were determined by differentiating Fourier polynomials of starch and sugar dynamics with respect to time. Hence, the Fourier polynomial balance models comprised three input functions, FP_input_ (Eqs. 1–3), and two balance equations, BE_1,2_ (Eqs. 4,5).

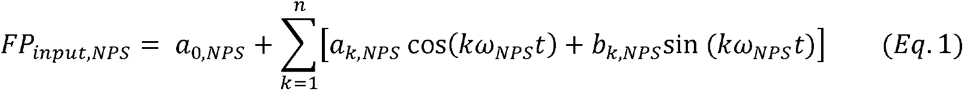

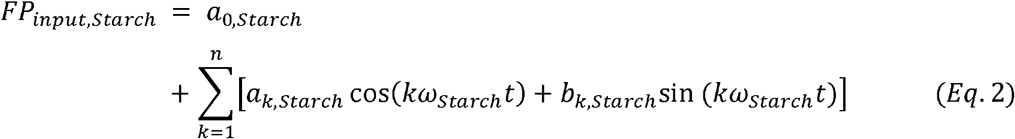

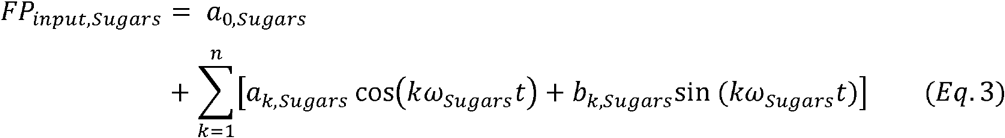

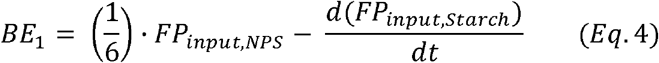

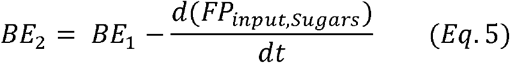

Here, a_k_ and b_k_ represent the Fourier coefficients for NPS, starch and sugar equations. ω is the fundamental frequency of the signal (ω = 2π/T, where T is the period). This Fourier polynomial-based balance equation model was applied to simulate dynamics of carbohydrate metabolism in plants of *Arabidopsis thaliana*, accession Columbia-0, under transient heat exposure. Additionally, NPS and carbohydrate dynamics were recorded and simulated in a starch-deficient mutant *pgm1* and a mutant with deficiency in sucrose biosynthesis capacity, *spsa1*. Coefficients of Fourier polynomials are provided in the supplements (Table S1).

### Fourier polynomials reflect dynamics of net CO_2_ assimilation rates

During the first 30-45 min of the light period, rates of net CO_2_ assimilation increased steeply in all genotypes and reached a first plateau at ~1250 μmol CO_2_ h^−1^ gDW^−1^ which was stable during the first half of the light period before it slightly increased until the end of the day at 22°C (Fig. 2, grey coloured lines). No significant difference was observed between genotypes, yet *spsa1* had slightly higher assimilation rates compared to Col-0 while rates of *pgm1* were slightly lower (Fig. 2 a, d, g). Temperature increase from 22°C to 32°C resulted in a drop of assimilation rates during the first hour of the treatment before the rates stabilized again and reached similar values than in the control (22°C) experiment (Fig. 2 a, d, g). At 32°C, starch deficient *pgm1* plants were most susceptible and mean values differed most from 22°C rates (Fig. 2 d). During the last 2 hours of the light period in which temperature was decreased to 22°C, all genotypes increased assimilation rates to control rates again. A similar scenario was observed within the 36°C experiment for *pgm1* and *spsa1* while Col-0 had significantly decreased assimilation rates during the last two hours of temperature treatment compared to the control experiment (Fig. 2 b). In *pgm1*, assimilation rates dropped significantly during the first 30 min of the recovery phase, i.e., between 6h and 6.5h, when temperature was decreased from 36°C to 22°C (Fig. 2 e). Such a significant recovery drop was also observed for both *pgm1* and *spsa1* mutants in the 40°C experiment, but not for Col-0 which showed again significantly decreased CO_2_ assimilation rates between the last two hours of the temperature treatment, i.e. between 4h and 6h of the light period (Fig.2 c, f, i). Experimentally determined mean values of CO_2_ assimilation rates and, by this, all described effects were covered by Fourier polynomials with R^2^ > 0.94 (exception: *pgm1*, 32°C, R^2^=0.8177).

### Transient heat exposure significantly affects dynamics of starch and soluble carbohydrates

The exposure to transient heat lead to a significant change in starch dynamics in Col-0 and *spsa1* (Fig. 3). Starch amount in *pgm1* was below the detection limit of the applied photometric detection method (Fig. 3 e-h). Starch concentration dropped significantly after transient heat exposure (6h) in comparison to control conditions in Col-0 and *spsa1* (p<0.001). This drop did not change significantly among the different temperatures. In the recovery phase after the heat shock (8h), plants increased their starch content depending on the temperature they were subjected to. Plants of both genotypes treated with 32°C increased their starch concentration by approximately 40% between 6h and 8h (Fig. 3 b, j), whereas plants treated with 36°C increased it by over 90% (Fig. 3 c, k). Col-0 subjected to the 36°C transient heat exposure was even able to reach the level of the control plants at the 8h time point.

**Figure 3.**
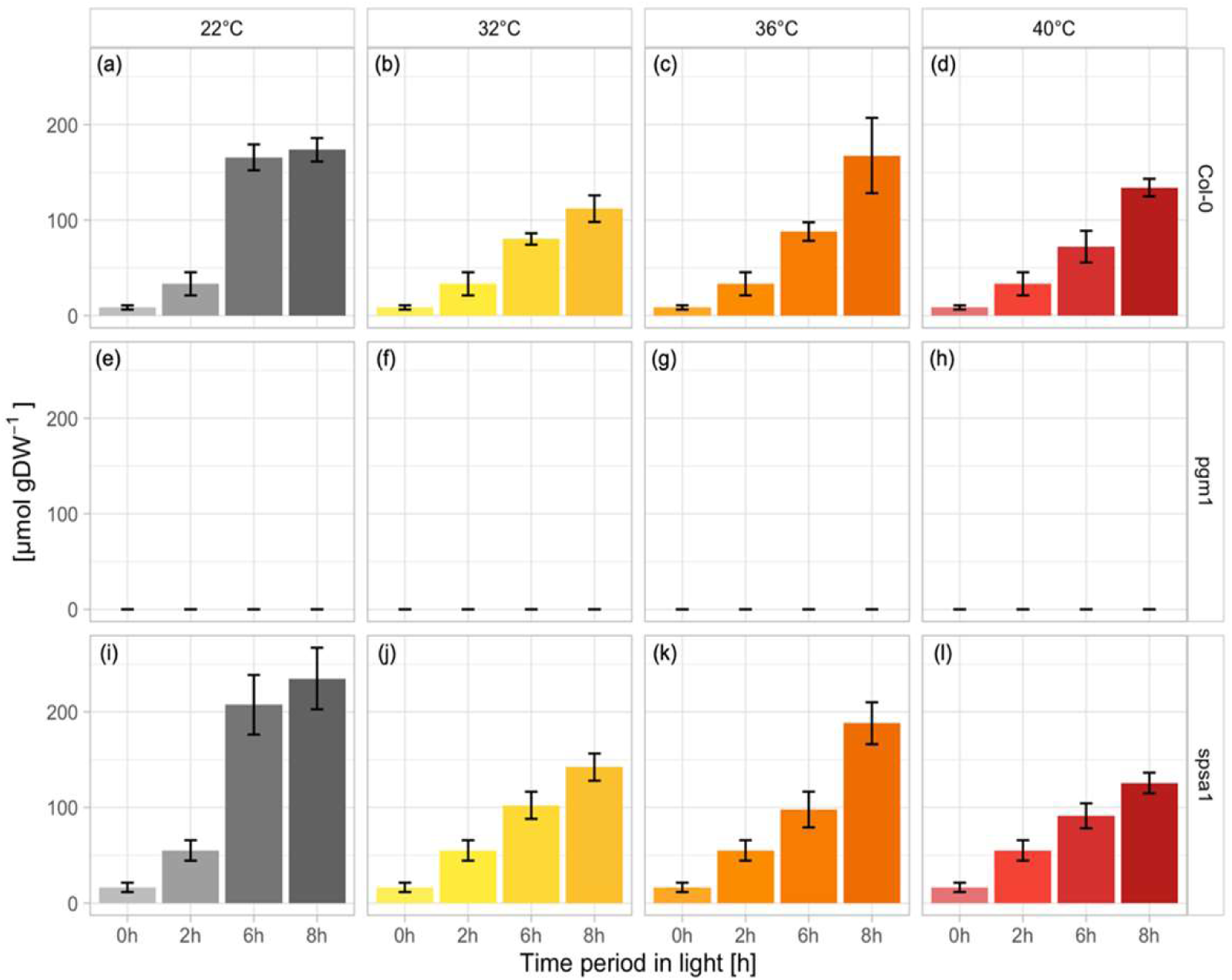
Starch amounts during short day transient heat exposure in glucose equivalents. Error bars represent mean values +/− SD of experimental data. (a) - (d): Col-0 (n≥5); (e) - (h): *pgm1* (n≥3); (i) - (l): *spsa1* (n≥5). Grey bars: 22°C experiment; yellow bars: 32°C experiment; orange bars: 36°C experiment; red bars: 40°C experiment. Temperature was set to 22°C between 0h-2h and 6h-8h. Temperature was transiently increased between 2h and 6h.

An effect of transient heat exposure on sucrose levels was only detectable for higher temperatures, i.e., within 36°C and 40°C experiments (Fig. 4). When subjected to 32°C of transient heat, no significant change in sucrose levels was detected in all genotypes (Fig. 4 b, f, j). In Col-0, only heat exposure of 40°C resulted in a significant increase in sucrose concentration at the 6h time point (p<0.001), but no further change was observed after the recovery phase at 8h (Fig. 4 d). Under all conditions, *pgm1* accumulated more sucrose over the course of the light phase compared to Col-0 and *spsa1* (Fig. 4 e-h). Nevertheless, heat treatment with 36°C and 40°C reduced the amount of sucrose in the *pgm1* plants almost significantly (p=0.05). In the recovery phase, the sucrose concentration in the heat treated *pgm1* plants returned to a level comparable to control conditions (Fig. 4 e-h). Due to high variance in the sucrose measurements of *spsa1* plants, there was no significant difference between Col-0 and *spsa1* in the control plants and the plants exposed to 32°C and 36°C of transient heat (Fig. 4 i-k). Only at 40°C, *spsa1* showed significantly higher sucrose levels than Col-0 at 40°C before and after the recovery phase (6h and 8h, p<0.001) or *spsa1* under control conditions (6h: p<0.004, 8h: p<0.01), whilst exhibiting very low variance (Fig. 4 l).

**Figure 4.**
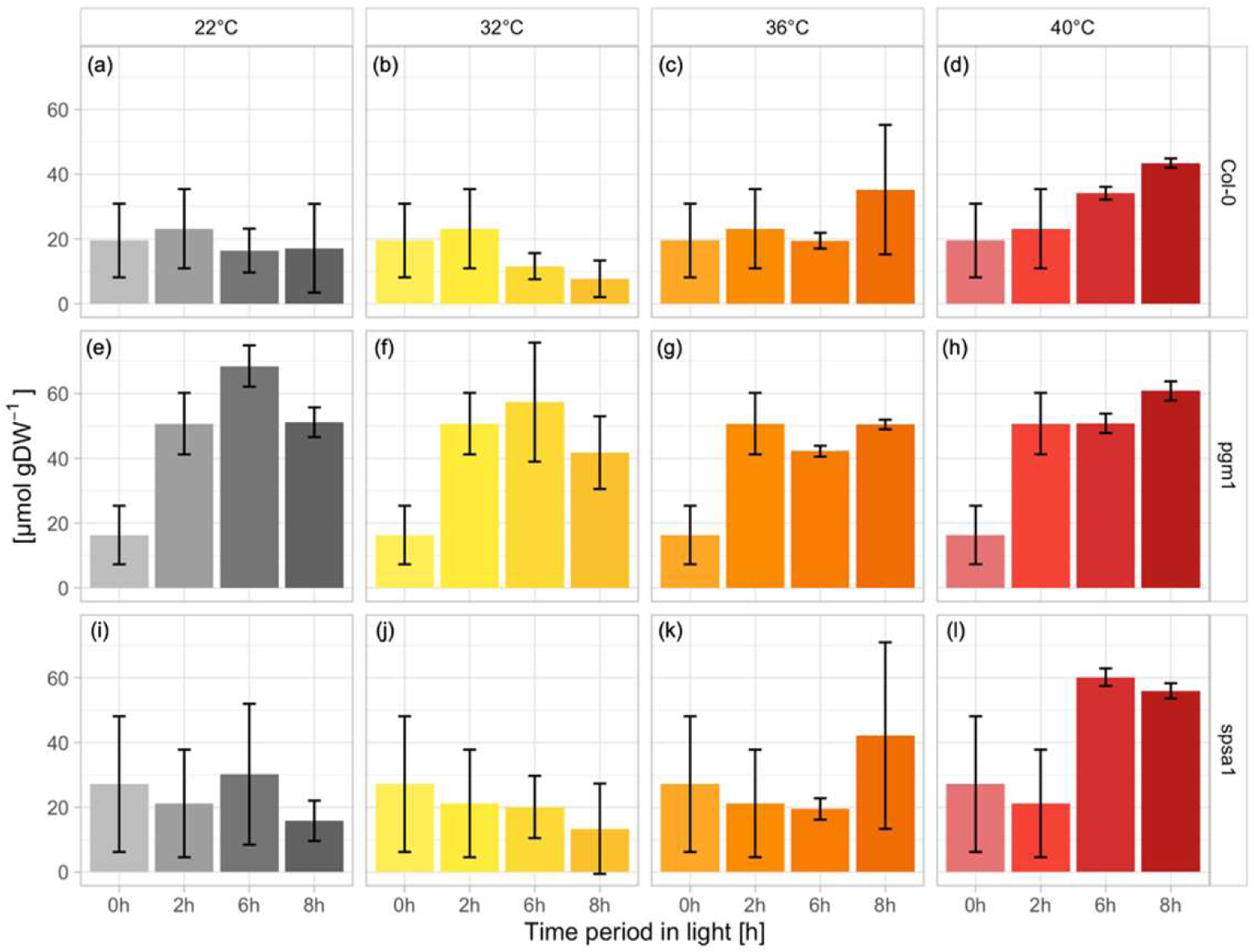
Sucrose concentrations during short day transient heat exposure. Error bars represent mean values +/− SD of experimental data. (a) - (d): Col-0 (n≥5); (e) - (h): *pgm1* (n≥3); (i) - (l): *spsa1* (n≥5). Grey bars: 22°C experiment; yellow bars: 32°C experiment; orange bars: 36°C experiment; red bars: 40°C experiment. Temperature was set to 22°C between 0h-2h and 6h-8h. Temperature was transiently increased between 2h and 6h.

In *pgm1*, glucose and fructose dynamics differed significantly from Col-0 (Fig. 5). Starting at equal hexose content at the start of the light period, the difference between *pgm1* and Col-0 and *spsa1* increased almost tenfold over the course of the day. Whilst *pgm1* increased hexoses steeply under control conditions, reaching a plateau at 6h, hexose concentrations in Col-0 and *spsa1* peaked at 2h, decreasing again afterwards.

**Figure 5.**
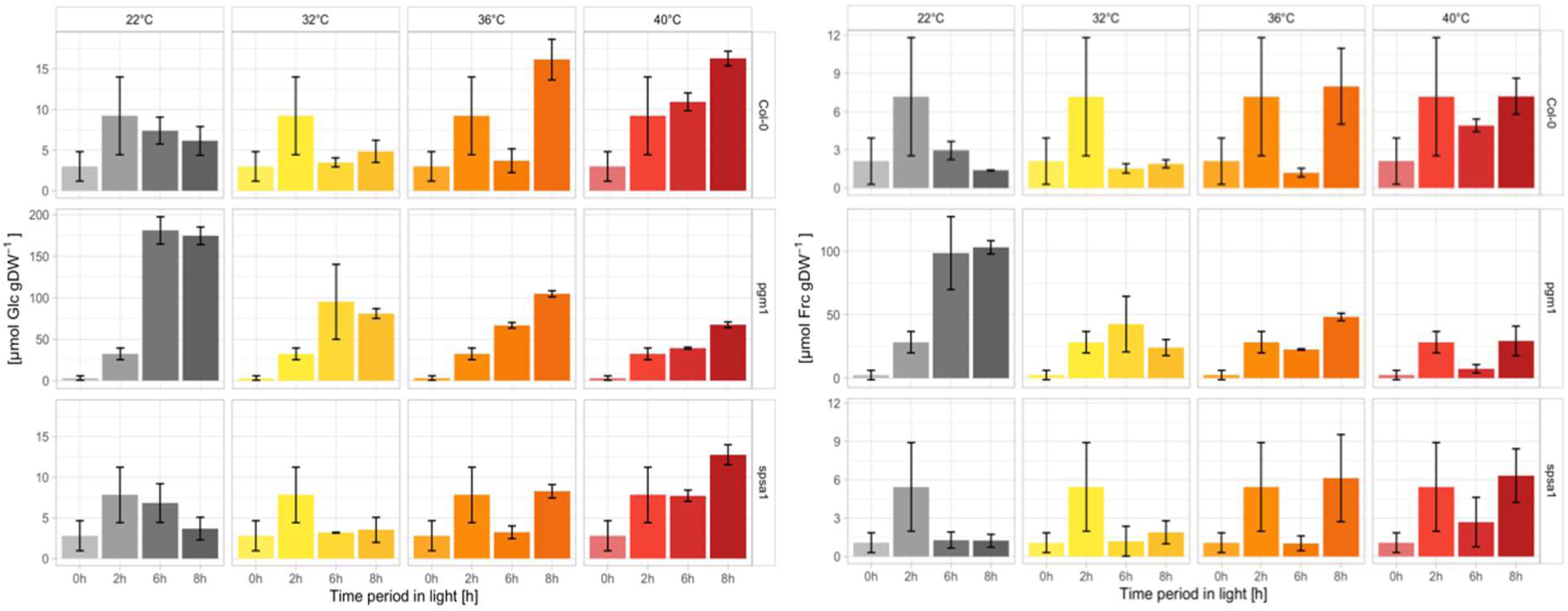
Hexose concentrations during short day transient heat exposure. Error bars represent mean values +/− SD of experimental data. Range of the y-axes differs for *pgm1* due to high difference in concentration. (a) Glucose concentrations; (b) Fructose concentrations. Top row: Col-0 (n≥5); middle row: *pgm1* (n≥3); bottom row: *spsa1* (n≥5). Grey bars: 22°C experiment; yellow bars: 32°C experiment; orange bars: 36°C experiment; red bars: 40°C experiment. Temperature was set to 22°C between 0h-2h and 6h-8h. Temperature was transiently increased between 2h and 6h.

Col-0 and *spsa1* showed a significant drop in glucose levels after heat exposure to 32°C and 36°C (p<0.001, Fig. 5 a). Within the 32°C experiment, glucose levels at 8h after recovery did not differ significantly from control conditions. Within the 36°C and 40°C experiments, however, the recovery phase after heat exposure resulted in a significant increase in glucose concentration compared to control conditions (p<0.001). Additionally, in Col-0 glucose levels were increasing above the level of control plants already during heat exposure to 40°C at 6h (p<0.001). Fructose dynamics in Col-0 were similar to the glucose dynamics in Col-0 (Fig. 5 b). In *spsa1*, however, fructose dynamics did not change significantly in response to transient heat exposure. The only differences were observable after the recovery phase at 8h in 36°C and 40°C. Here, a significantly higher fructose content could be measured compared to 22°C (p<0.001). In *pgm1,* hexose levels decreased significantly after temperature treatment (6h, p<0.001), with the lowest values being reached at 40°C. After recovery (8h), hexose levels did not change significantly from the 6h time point in the 32°C plants. After exposure to 36°C and 40°C, however, hexose levels increased significantly from 6h to 8h (p<0.001).

### Numerical differentiation and integration of carbon balance equations reveals genotype-dependent system fluctuations due to transient heat exposure

To reveal how dynamics of carbon balance equations, which combine net photosynthesis, starch (BE_1_) and sugar metabolism (BE_2_), are affected by transient heat exposure, derivatives were built with respect to time (Fig. 6). In Col-0 and *spsa1*, increasing temperature resulted in decreased fluctuation of BE_1_ and BE_2_ derivatives (Fig. 6 a, b, e, f). Particularly in 36°C and 40°C experiments, oscillations of derivatives were significantly damped during the second half of the temperature stress period, i.e. between 4h and 6h of the light period. In Col-0, this dampening effect was strongest at 40°C, in *spsa1* at 36°C. In both genotypes, the 32°C experiment had only little effect on derivatives, i.e., on absolute changing rates of Fourier polynomials (yellow lines in Fig. 6 a, b, e, f). In *pgm1*, fluctuation of derivatives increased for both BE_1_ and BE_2_ under 32°C and 36°C compared to 22°C (Fig. 6 c, d; grey (22°C), yellow (32°C) and orange (36°C) lines). Within the 40°C experiment, fluctuations of derivative fluctuations were slightly damped in *pgm1*, and rates became negative in BE_2_ during the second half of heat exposure, i.e., between 4h and 6h before they increased again until 7h in the light period (Fig. 6 d; red lines). In addition to derivative functions which revealed the absolute changing rates of balance equations, fundamental frequencies of Fourier polynomials of BE_1_ and BE_2_ further suggested a differential effect of high temperature on genotypes’ carbon balances (Fig. 7). In Col-0, frequencies of BE_1_ and BE_2_ under 32°C and 36°C were almost double as high as in the 22°C experiment before they dropped in the 40°C experiment. This decrease was more emphasized in BE_2_ than in BE_1_ indicating a contribution of sugar dynamics. In *pgm1*, frequencies of BE_1_ constantly increased with temperature in experiments (Fig. 7 a). Frequencies of BE_2_ peaked under 32°C and, finally, were lower under 40°C than under 22°C (Fig. 7 b). In *spsa1*, dynamics of BE_1_ and BE_2_ frequencies across experiments were similar, yet more pronounced in BE_2_. In contrast to Col-0 and *pgm1*, lowest frequency of both balance equations was observed for the 36°C experiment.

**Figure 6.**
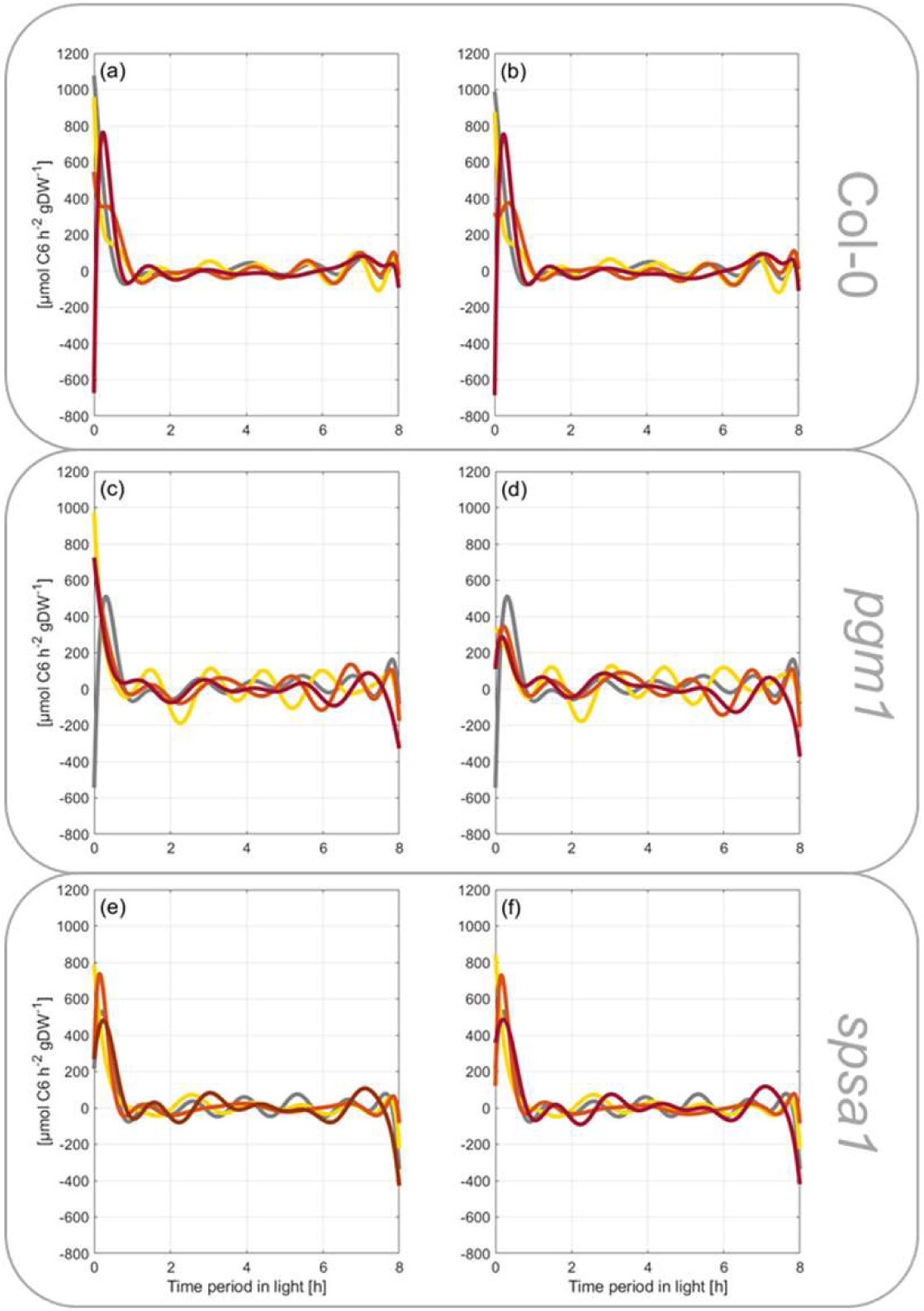
Derivatives of carbon balance equations with respect to time. Derivatives of balance equations were built for the experiments “22°C” (control; grey lines), “32°C” (yellow lines), “36°C” (orange lines) and “40°C” (red lines). Left panel represents derivatives of balance equation 1 (BE_1_) for Col-0 (a), *pgm1* (c) and *spsa1* (e). Right panel represents derivatives of balance equation 2 (BE_2_) for Col-0 (b), *pgm1* (d) and *spsa1* (f).

**Figure 7.**
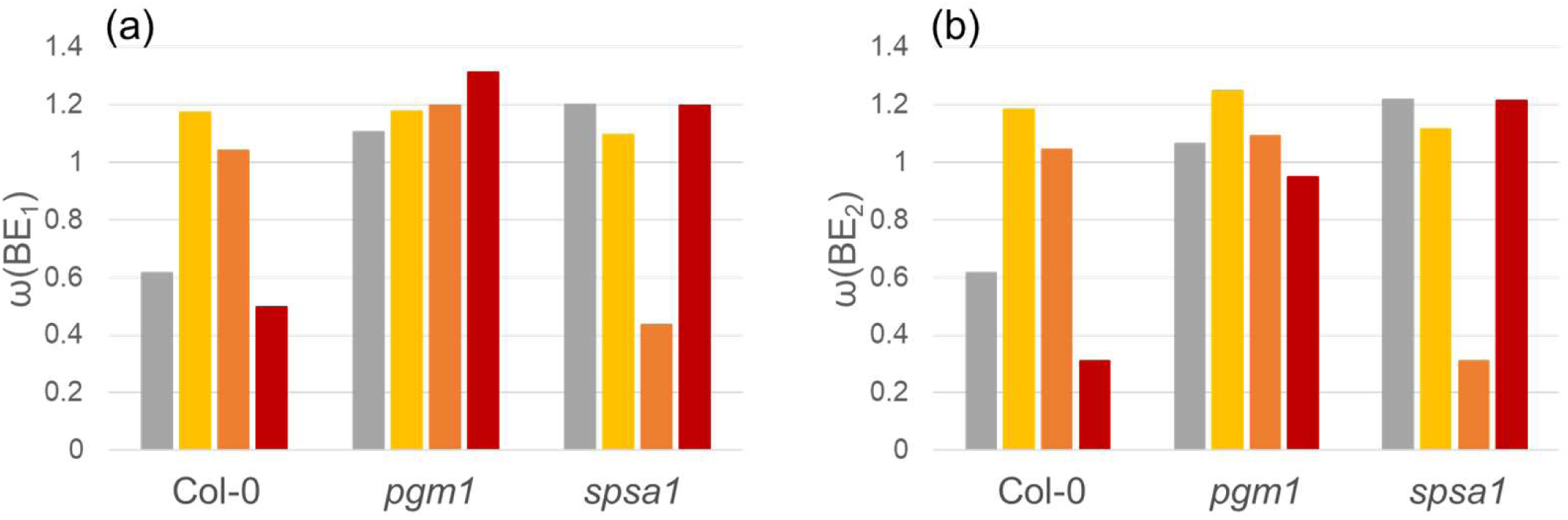
Fundamental frequencies of Fourier polynomials of BE_1_ (a) and BE_2_ (b). Frequencies were calculated as the ratio of (2π/T) where T represents periods. Bar colour indicates different experiments. Grey bars: 22°C experiment; yellow bars: 32°C experiment; orange bars: 36°C; red bars: 40°C experiment.

Numerical integration of NPS rates over time period in light revealed a decreased amount of assimilated carbon due to heat exposure in Col-0 (Fig. 8 a) and *pgm1* (Fig. 8 d). A detailed summary of numerical values of integrals is provided in the supplements (Table S2). In Col-0, transient heat effects on NPS rates became strongest after 2h of temperature treatment, i.e. after 4h in the light period. Between 4h and 6h, particularly the amount of carbon assimilated at 36°C and 40°C deviated clearly from the 22°C experiment (Fig. 8 a). In *pgm1*, this effect was observed two hours earlier, i.e., during the first two hours of heat treatment between 2h and 4h in the light period (Fig. 8 d). Surprisingly, however, the 40°C effect on carbon assimilation was not as strong as observed for 32°C and 36°C. In *spsa1*, carbon assimilation under heat was most robust and similar to control conditions, i.e., 22°C (Fig. 8 g).

**Figure 8.**
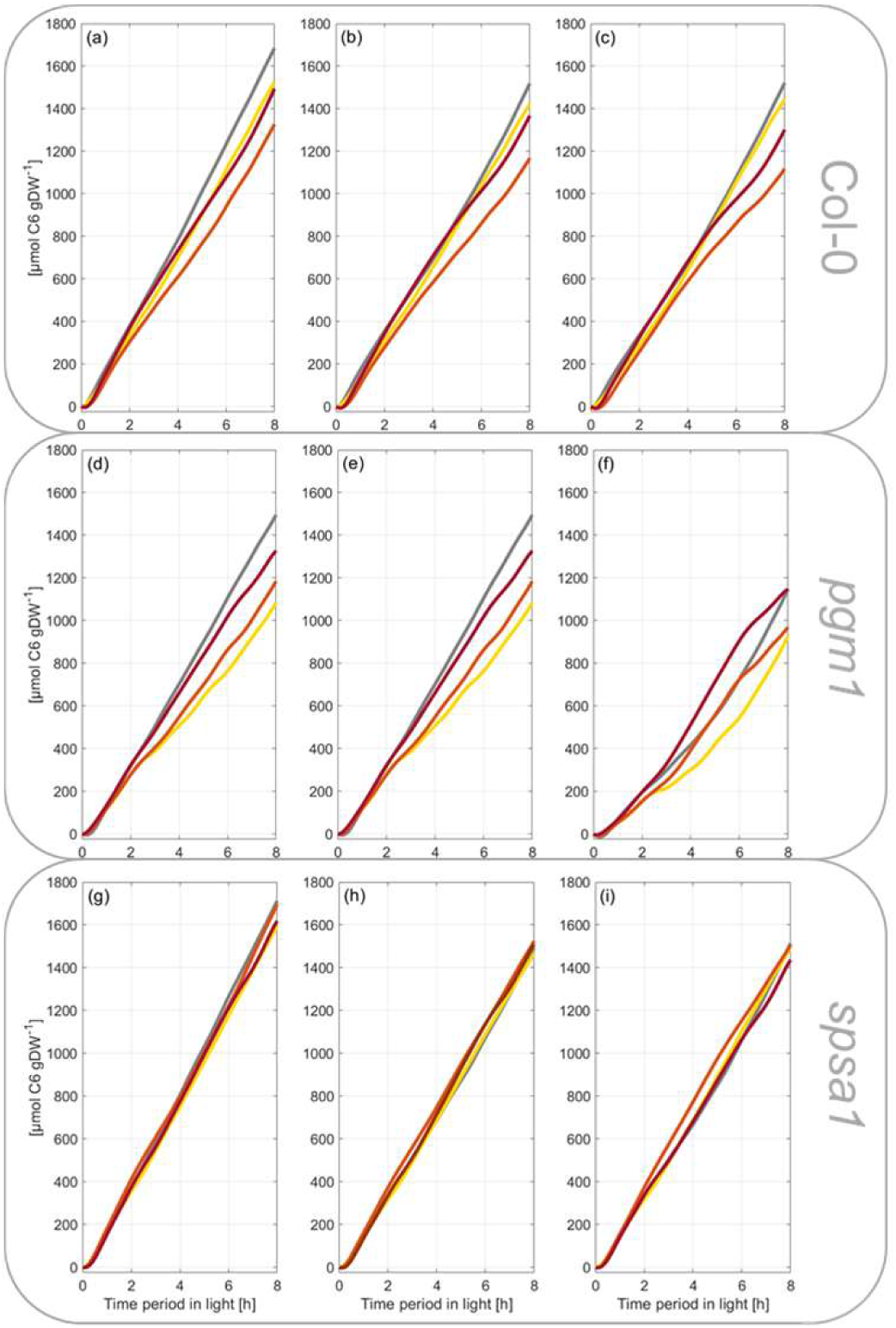
Integrals of carbon balance rates during transient heat exposure. NPS rates and rates derived from BE_1_ and BE_2_ were integrated over time to reveal the total sum of net carbon gain during the light period. **(a) – (c)** : Col-0 integrals of NPS rates **(a)**, BE_1_ **(b)** and BE_2_ **(c)**. **(d) – (f)** : *pgm1* integrals of NPS rates **(d)**, BE_1_ **(e)** and BE_2_ **(f)**. **(g) – (i)**: *spsa1* integrals of NPS rates **(g)**, BE_1_ **(h)** and BE_2_ **(i)**. Numerical values of integrals are provided in the supplements (Table S2).

Integrals of BE_1_, which in addition to NPS rates also considered starch dynamics, revealed that starch dynamics in Col-0 were adjusted proportionally to affected NPS rates during transient heat exposure (Fig. 8 b). In particular, integrals of 32°C and 40°C experiments became similar to the control experiment (22°C). Due to starch deficiency, this effect was not observed in *pgm1* (Fig. 8 e) while heat exposure resulted in larger integrals of BE_1_ in *spsa1* (Fig. 8 h). These heat-induced effects became more pronounced in BE_2_ integrals which additionally considered net carbon flux into soluble sugar biosynthesis. In Col-0, discrepancy of integrals between heat and control experiments was minimized during the first half of the light period, i.e., within the first two hours of heat exposure (Fig. 8 c). During the second half of the light period, discrepancy increased for 36°C and 40°C experiments. In *pgm1*, net carbon flux into sugar biosynthesis was reduced in a temperature-dependent manner which resulted in an (over-)compensation of reduced CO_2_ assimilation rates under 36°C and 40°C (Fig. 7 f). Also, in *spsa1* integrals of BE_2_ increased under transient heat but this effect was less pronounced than in *pgm1* and Col-0 (Fig. 8 i).

During the recovery phase, in which temperature was set to 22°C again (6h ➔ 8h of light period), changes in starch and sugar dynamics became obvious in integrals of BE_1_ and BE_2_ for all genotypes. In Col-0 this effect was most pronounced within 36°C and 40°C experiments (Fig. 8 b, c). In this phase, curves of integrals showed an inflection point directing the curve of integrals towards the control samples. In summary, this indicated reversibility of temperature-induced metabolic effects.

## Discussion

In temperate regions, plants are frequently exposed to a changing temperature regime, and these changes might occur over short and relatively long time scales. For example, temperature typically changes between day and night, and beyond, temperature might also change transiently within the diurnal light and dark period. While temperature acclimation of plants typically can be observed after days of exposure to non-lethal cold or heat (Atanasov *et al.*, 2020; Garcia-Molina *et al.*, 2020), transient temperature changes and plant stress response occurs within minutes or hours. Interestingly, *Arabidopsis thaliana* was found to memorize already 5 minutes of heat stress which indicates a tightly regulated molecular network involved in heat stress response (Oyoshi *et al.*, 2020). High temperature, e.g., between 35°C – 40°C, is well known to result in a reduced rate of photosynthesis (Sharkey, 2005) which has also been observed in the present study. While in Col-0 and *spsa1*, 32°C resulted in only slightly decreased NPS rates, higher temperatures of 36°C and 40°C resulted in a significantly decreased NPS rate during the second half of the heat exposure period only in Col-0 (see Fig. 2). As previously summarised, a decreased NPS rate is not due to photosystem damage, but rather due to rubisco deactivation (Sharkey, 2005). Consistent with previous findings which show a decreased rubisco activation at leaf temperature >35°C (Crafts-Brandner and Salvucci, 2000), the effect of 32°C on NPS rates was much less significant than at 36°C and 40°C. In *spsa1*, however, NPS rates were less affected by heat than in Col-0 which might have several reasons. First, compared to Col-0, *spsa1* might have had a reduced rate of photorespiration and/or mitochondrial respiration during heat exposure. While it remains speculation from our study, a higher starch accumulation rate in *spsa1* might result in a lowered respiration rate under heat because carbon equivalents can be fixed more efficiently. The observation of a destabilized NPS rate in starchless *pgm1* plants would support the stabilizing role of starch biosynthesis under transient heat exposure. However, previous reports under ambient conditions have shown that SPS knockout mutants have rather enhanced than lowered dark respiration rates which does not directly support this hypothesis (Bahaji *et al.*, 2015). Another explanation might be a secondary effect of the *spsa1* mutation on rubisco and/or rubisco activase which, to our knowledge, has not been shown in current literature but which needs to be excluded in future studies.

### Fructose concentration profiles are more conserved than glucose and sucrose profiles during transient heat exposure

While sucrose and glucose metabolism showed a dynamic and differential accumulation profile between 32°C, 36°C and 40°C experiments, dynamics of fructose concentrations were most conserved across all temperature treatments and, remarkably, also across genotypes (see Fig. 5). An initial accumulation within the first 2h of the light period was followed by a significant decrease until the end of heat exposure and an accumulation during the recovery phase between 6h and 8h of the light period. Only *pgm1* showed a differential pattern at 22°C and 32°C but became similar in its fructose profile to Col-0 and *spsa1* under 36°C and 40°C. In mature *Arabidopsis* leaves, fructose levels are significantly affected by invertases which catalyze hydrolysis of sucrose and release free hexoses (Wan *et al.*, 2018), and by fructokinase catalyzing ATP-dependent phosphorylation which yields fructose-6-phosphate (Claeyssen and Rivoal, 2007). As fructose and glucose profiles differed in the present study, this cannot (solely) be explained by invertase reactions which release equimolar concentrations of both hexoses. However, differential regulation of hexokinase and fructokinase could explain the different hexose profiles. Fructokinase yields the direct substrate for glycolysis, TCA cycle and mitochondrial respiration. In a previous study which analysed transcript levels in *Arabidopsis thaliana* under combined drought and heat stress found increased transcripts for both hexokinase and fructokinase (Rizhsky *et al.*, 2004). Although the experimental design differed significantly from the present study, together with other findings this suggests a central role of hexose phosphorylation in heat stress response and acclimation (Atanasov *et al.*, 2020). As leaf respiration rates typically increase under elevated temperature (Dusenge *et al.*, 2019), observed consistent fructose dynamics might be due to a relatively high rate of glycolytic consumption under transient heat exposure.

### Data integration by Fourier polynomials reveals dynamic properties of plant carbon metabolism in a changing environment

Integrating net CO_2_ assimilation rates with starch and sugar turnover allows for balancing of the central carbohydrate metabolism. In this context, Fourier polynomials support the functional and time-continuous estimation of dynamics of metabolism. Integration and differentiation of Fourier polynomials is straight forward, and, at the same time, provides a comprehensive mathematical framework which is applied in diverse fields of natural sciences and engineering (Li *et al.*, 2018; Plonka *et al.*, 2018; Wang *et al.*, 1997). As described in the block diagram model (see Fig. 1), metabolic dynamics were simulated by addition of Fourier polynomials comprising input functions (NPS rates) and consuming functions (starch and sugar dynamics). With such a design, dynamics of plant carbon balancing become traceable without the need for application of composed spline functions. Further, properties of Fourier polynomials can reveal further insight into metabolic regulation and consequences of environmental changes. For example, in the present study both amplitude and frequency of derivatives of balancing equations differed with regard to genotype and environment. A different pattern was observed in Col-0 and *spsa1* than in *pgm1*, indicating that the starchless mutant has a less buffered metabolic response towards heat stress than both other genotypes. This was supported by the comparison of fundamental frequencies of BE_1_ and BE_2_ Fourier polynomials. Here, a genotype-specific pattern was observed which reflected the impact of starch deficiency in *pgm1* and comparatively high metabolic dynamics in *spsa1* within the 40°C experiment (see Fig. 4 and 5). Thus, summarizing effects of transient heat on NPS rates and carbohydrate metabolism resulted in characteristic Fourier polynomials which enabled the discrimination of genotypes by their derivatives and fundamental frequencies. This discrimination was rather enabled by the dynamics of frequencies across different temperature experiments than their absolute value.

Fourier analysis and spectra of frequencies have been applied before in a different context, e.g., to analyse gene-expression data time-series data (Dong *et al.*, 2011). These authors coupled Fourier analysis to supervised learning algorithms to discriminate between housekeeping genes and non-housekeeping genes in HeLa cells. This example provides evidence for the suitability of Fourier analysis to be combined with machine learning algorithms which is of particular interest for large-scale data sets. However, also data sets with only a relatively low number of variables may need mathematical functions for quantitative analysis and integration, e.g., as shown in the present study. This is due to the need for combining dynamics of variables rather than steady-state values under one condition. Here, Col-0, *pgm1* and *spsa1* could successfully be discriminated by the dynamics of fundamental frequencies of Fourier polynomials across different experiments rather than by one absolute value of a frequency. This observation further emphasizes the need for a functional mathematical description of experimentally observed system dynamics because underlying attributes, e.g., monotonicity or curvature, can be derived from such a description. These attributes provide valuable information about system properties like stability or predictability which need to be essentially addressed for predictive modelling (Angeli, 2013; Sontag, 2007). Conclusively, Fourier analysis provides a mathematical approach which can essentially support nonlinear modelling of metabolism, and which might, in future studies, even serve as a mathematical framework to connect oscillations in metabolism with quantum theory (Iotti *et al.*, 2010; Jaffe *et al.*, 2020).

## Supplementary data

**Figure S1.** SPS maximum activity under substrate saturation in Col-0 and *spsa1*.

**Figure S2.** Schematic representation of the experimental design used in this study.

**Figure S3.** Summary of actual growth cabinet parameters during sampling period.

**Table S1.** Fourier polynomials and balance equations.

**Table S2.** Results of numerical integration of net carbon assimilation.

## Acknowledgements

We thank the members of Plant Evolutionary Cell Biology and Plant Development at LMU for fruitful discussions. We especially thank Laura Schröder, Anastasia Kitashova and Lisa Fürtauer for lab support.

## Author contributions

All authors performed experiments. CS and TN performed statistics and modelling and wrote the paper. All authors approved the paper.

## Data availability statement

The datasets used and/or analysed during the current study are provided in the manuscript and supplements.

## References

Angeli D. 2013. Monotone Systems in Biology. In: Baillieul J, Samad T, eds. Encyclopedia of Systems and Control. London: Springer London, 1–9.

Atanasov V, Fürtauer L, Nägele T. 2020. Indications for a central role of hexokinase activity in natural variation of heat acclimation in *Arabidopsis thaliana*. Plants 9, 819.

Bahaji A, Baroja-Fernández E, Ricarte-Bermejo A, Sánchez-López ÁM, Muñoz FJ, Romero JM, Ruiz MT, Baslam M, Almagro G, Sesma MT, Pozueta-Romero J. 2015. Characterization of multiple SPS knockout mutants reveals redundant functions of the four Arabidopsis sucrose phosphate synthase isoforms in plant viability, and strongly indicates that enhanced respiration and accelerated starch turnover can alleviate the blockage of sucrose biosynthesis. Plant Science 238, 135–147.

Basler G, Fernie Alisdair R, Nikoloski Z. 2018. Advances in metabolic flux analysis toward genome-scale profiling of higher organisms. Bioscience Reports 38.

Claeyssen E, Rivoal J. 2007. Isozymes of plant hexokinase: occurrence, properties and functions. Phytochemistry 68, 709–731.

Costa RS, Hartmann A, Vinga S. 2016. Kinetic modeling of cell metabolism for microbial production. Journal of Biotechnology 219, 126–141.

Crafts-Brandner SJ, Salvucci ME. 2000. Rubisco activase constrains the photosynthetic potential of leaves at high temperature and CO_2_. Proceedings of the National Academy of Sciences 97, 13430–13435.

Crown SB, Long CP, Antoniewicz MR. 2016. Optimal tracers for parallel labeling experiments and 13C metabolic flux analysis: A new precision and synergy scoring system. Metabolic Engineering 38, 10–18.

Dong B, Zhang P, Chen X, Liu L, Wang Y, He S, Chen R. 2011. Predicting housekeeping genes based on Fourier analysis. PLoS ONE 6, e21012.

Dusenge ME, Duarte AG, Way DA. 2019. Plant carbon metabolism and climate change: elevated CO2 and temperature impacts on photosynthesis, photorespiration and respiration. New Phytologist 221, 32–49.

Espinoza C, Degenkolbe T, Caldana C, Zuther E, Leisse A, Willmitzer L, Hincha DK, Hannah MA, Harmon F. 2010. Interaction with Diurnal and Circadian Regulation Results in Dynamic Metabolic and Transcriptional Changes during Cold Acclimation in Arabidopsis. PLoS ONE 5, e14101.

Feldman-Salit A, Veith N, Wirtz M, Hell R, Kummer U. 2019. Distribution of control in the sulfur assimilation in Arabidopsis thaliana depends on environmental conditions. New Phytologist 222, 1392–1404.

Garcia-Molina A, Kleine T, Schneider K, Mühlhaus T, Lehmann M, Leister D. 2020. Translational components contribute to acclimation responses to high light, heat, and cold in Arabidopsis. iScience 23, 101331.

Gutenkunst RN, Waterfall JJ, Casey FP, Brown KS, Myers CR, Sethna JP. 2007. Universally sloppy parameter sensitivities in systems biology models. PLoS Computational Biology 3, 1871–1878.

Iotti S, Borsari M, Bendahan D. 2010. Oscillations in energy metabolism. Biochimica et Biophysica Acta (BBA) - Bioenergetics 1797, 1353–1361.

Jaffe A, Jiang C, Liu Z, Ren Y, Wu J. 2020. Quantum Fourier analysis. Proceedings of the National Academy of Sciences 117, 10715–10720.

Kitashova A, Schneider K, Fürtauer L, Schröder L, Scheibenbogen T, Fürtauer S, Nägele T. 2021. Impaired chloroplast positioning affects photosynthetic capacity and regulation of the central carbohydrate metabolism during cold acclimation. Photosynthesis Research 147, 49–60.

Li J, Liu P, Yu W, Cheng X. 2018. The morphing of geographical features by Fourier transformation. PLoS ONE 13, e0191136.

Lopatkin AJ, Collins JJ. 2020. Predictive biology: modelling, understanding and harnessing microbial complexity. Nature Reviews Microbiology 18, 507–520.

Mochão H, Barahona P, Costa RS. 2020. KiMoSys 2.0: an upgraded database for submitting, storing and accessing experimental data for kinetic modeling. Database 2020.

Nägele T, Fürtauer L, Nagler M, Weiszmann J, Weckwerth W. 2016. A strategy for functional interpretation of metabolomic time series data in context of metabolic network information. Frontiers in Molecular Biosciences 3, 6.

Nägele T, Heyer AG. 2013. Approximating subcellular organisation of carbohydrate metabolism during cold acclimation in different natural accessions ofArabidopsis thaliana. New Phytologist 198, 777–787.

Oyoshi K, Katano K, Yunose M, Suzuki N. 2020. Memory of 5-min heat stress in Arabidopsis thaliana. Plant Signaling and Behavior, 1778919.

Plonka G, Potts D, Steidl G, Tasche M. 2018. Numerical Fourier Analysis: Springer.

R Core Team. 2019. R: A language and environment for statistical computing. R Foundation for Statistical Computing, Vienna, Austria.

Ramos MPM, Ribeiro C, Soares AJ. 2019. A kinetic model of T cell autoreactivity in autoimmune diseases. Journal of Mathematical Biology 79, 2005–2031.

Rizhsky L, Liang H, Shuman J, Shulaev V, Davletova S, Mittler R. 2004. When defense pathways collide. The response of Arabidopsis to a combination of drought and heat stress. Plant Physiology 134, 1683–1696.

Rohwer JM. 2012. Kinetic modelling of plant metabolic pathways. Journal of Experimental Botany 63, 2275–2292.

Sajitz-Hermstein M, Töpfer N, Kleessen S, Fernie AR, Nikoloski Z. 2016. iReMet-flux: constraint-based approach for integrating relative metabolite levels into a stoichiometric metabolic models. Bioinformatics 32, i755–i762.

Schaber J, Liebermeister W, Klipp E. 2009. Nested uncertainties in biochemical models. IET Systems Biology 3, 1–9.

Sharkey TD. 2005. Effects of moderate heat stress on photosynthesis: importance of thylakoid reactions, rubisco deactivation, reactive oxygen species, and thermotolerance provided by isoprene. Plant, Cell and Environment 28, 269–277.

Sontag ED. 2007. Monotone and near-monotone biochemical networks. Systems and Synthetic Biology 1, 59–87.

Wan H, Wu L, Yang Y, Zhou G, Ruan YL. 2018. Evolution of sucrose metabolism: the dichotomy of invertases and beyond. Trends in Plant Science 23, 163–177.

Wang L, Bert JL, Okazawa M, Paré PD, Pinder KL. 1997. Fast Fourier transform analysis of dynamic data: sine wave stress - strain analysis of biological tissue. Physics in Medicine and Biology 42, 537–547.

Weiszmann J, Fürtauer L, Weckwerth W, Nägele T. 2018. Vacuolar sucrose cleavage prevents limitation of cytosolic carbohydrate metabolism and stabilizes photosynthesis under abiotic stress. The FEBS Journal 285, 4082–4098.

