## Supplementary material for "Integration of plant carbohydrate dynamics by Fourier polynomials": Figure S1

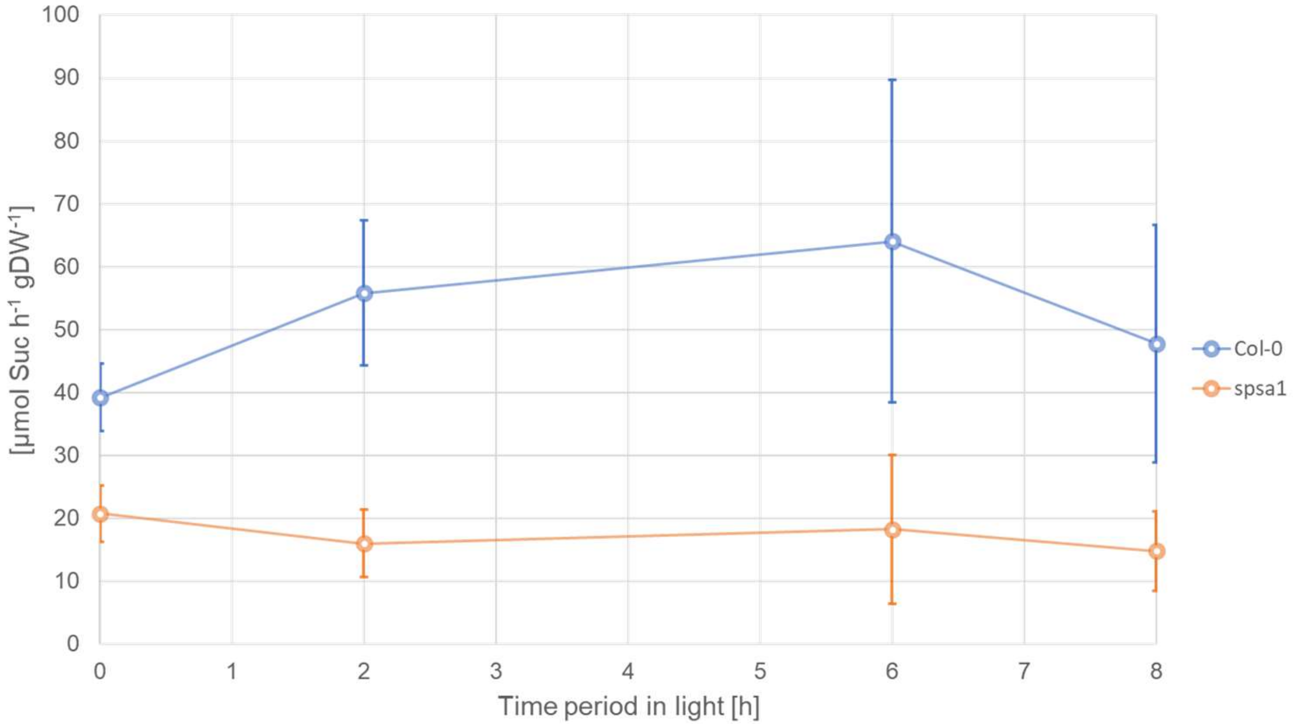


**Figure S1. SPS maximum activity under substrate saturation in Col-0 and *spsa1*.** Blue line: Col-0, orange line: *spsa1*. Errorbars represent means +/- SD (n = 5).
