## Supplementary material for "Integration of plant carbohydrate dynamics by Fourier polynomials": Figure S2

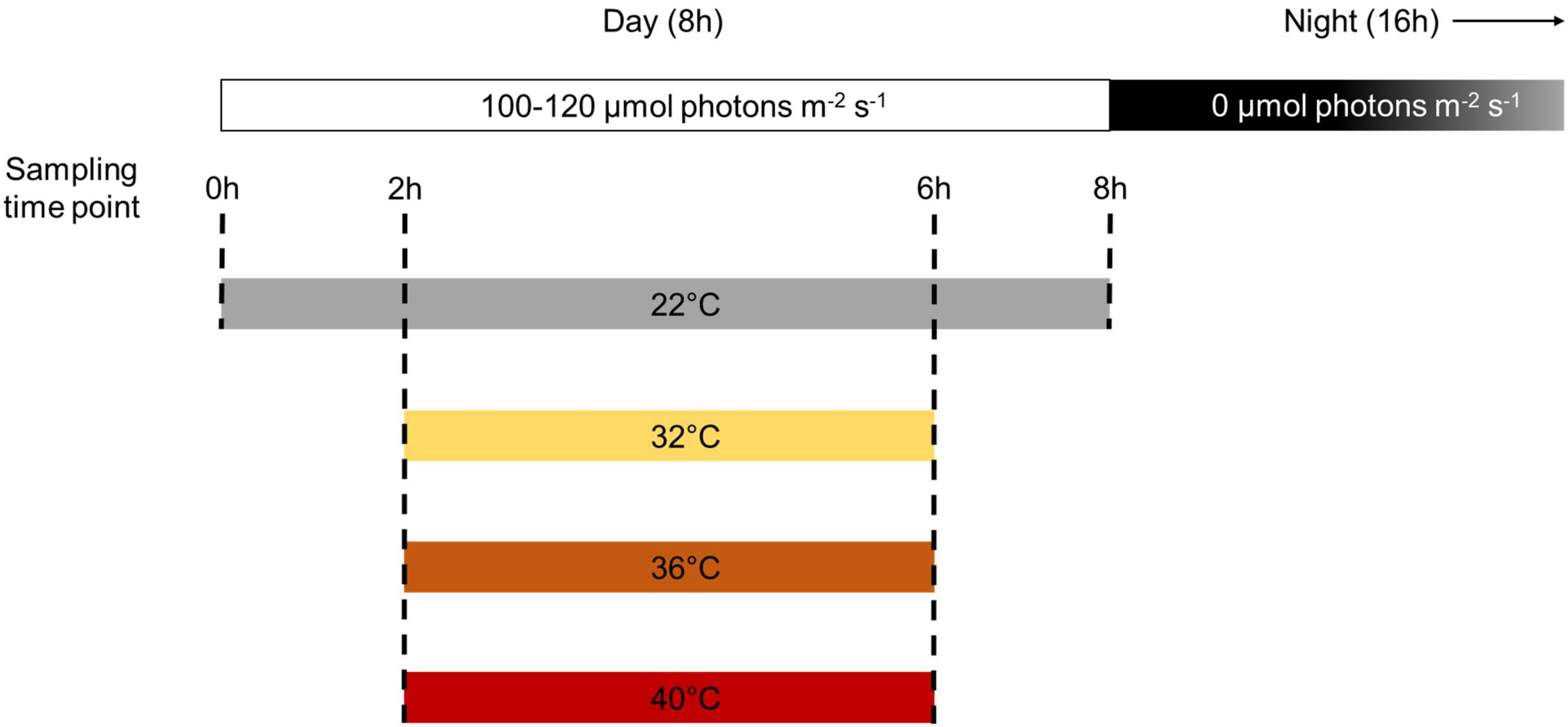


**Figure S2. Schematic representation of the experimental design used in this study.** Sampling time points are indicated by dashed lines. Experiments were entitled based on the temperature setpoint of the growth cabinet (“22°C” or “control”, “32°C”, “36°C” and “40°C”) between 2h and 6h in the light period. However, actual temperature gradients measured during the experiments deviated and are summarized in Figure S3.
