## Supplementary material for "Integration of plant carbohydrate dynamics by Fourier polynomials": Figure S3

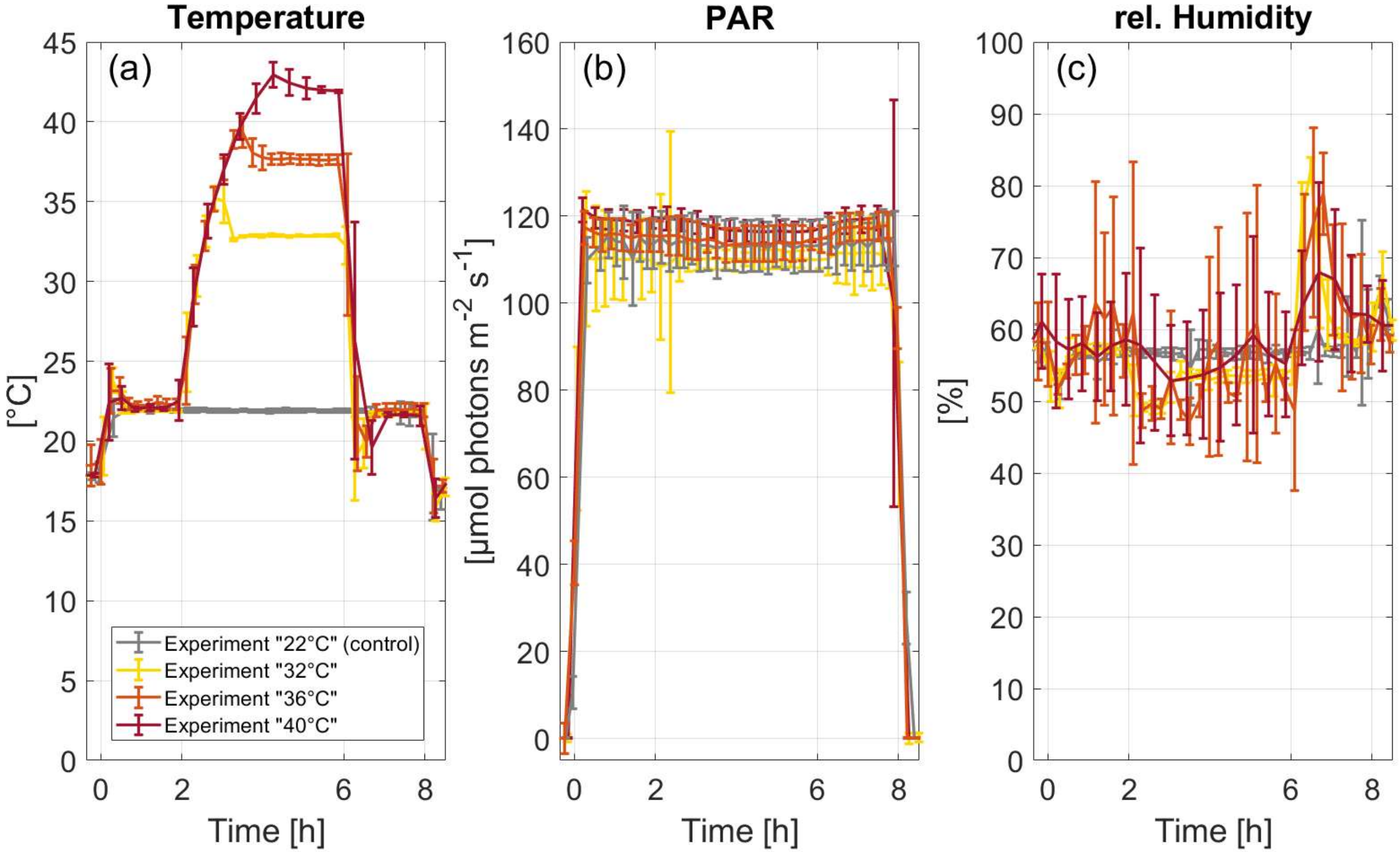


**Figure S3. Summary of actual growth cabinet parameters during sampling period.** Parameters were recorded inside the measurement head during net photosynthesis measurements. Errorbars represent means +/- SD of n = 3 measurements for each experiment.
